# Towards stability of dynamic FC estimates in neuroimaging and electrophysiology: solutions and limits

**DOI:** 10.1101/2023.01.18.524539

**Authors:** Sonsoles Alonso, Diego Vidaurre

## Abstract

Time-varying functional connectivity methods are used to map the spatiotemporal organization of brain activity. However, their estimation can be unstable, in the sense that different runs of the inference may yield different solutions. But to draw meaningful relations to behaviour, estimates must be robust and reproducible. Here, we propose two solutions using the Hidden Markov Model (HMM) as a descriptive model of time-varying FC. The first, best-ranked HMM, involves running the inference multiple times and selecting the best model based on a quantitative measure combining fitness and model complexity. The second, hierarchical clustered HMM, generates stable aggregated state timeseries by applying hierarchical clustering to the state timeseries obtained from multiple runs. Experimental results on fMRI and MEG data demonstrate that these approaches substantially improve the stability of time-varying FC estimations. Overall, hierarchical clustered HMM is preferred when the inference variability is high, while the best-ranked HMM performs better otherwise.

## Introduction

An important aspect of the brain’s functional architecture is how different regions are brought together into functional networks, and how these are dynamically organized at different spatial and temporal scales (Laughlin and Sejnowski, 2003)□. One of the most widely used metrics to map these functional relationships is functional connectivity (FC), a measure of the statistical dependencies between pairs of brain regions (Friston, 1994)□. More recently, the exploration of the temporal properties of these interdependencies has revealed meaningful within-session fluctuations in FC, both from functional magnetic resonance imaging (fMRI; Fornito and Bullmore, 2010; Karapanagiotidis et al., 2020; Liégeois et al., 2019; Lurie et al., 2020; Vidaurre et al., 2021, 2018, 2017; Xie et al., 2018)□□ and electrophysiology (de Pasquale et al., 2010, 2012; Baker et al., 2014; Vidaurre et al., 2018).

Unfortunately, comparing results across studies can be challenging due to the variety of analytical tools for estimating time-varying FC and their inherent limitations (Dafflon et al., 2022). One common method for estimating time-varying FC is the Hidden Markov Model (HMM; Vidaurre et al., 2017), where an optimization process aims to determine the best model parameters that fit the observed timeseries data. However, due to the random initialization of the optimization algorithm, different runs of the algorithm may produce different results even on the same data. Consequently, the estimation of time-varying FC can be unstable, and this instability may depend on factors such as the amount of data and the complexity of the model (Vidaurre et al., 2019)⍰. This is the case also for simpler approaches, such as Independent Component Analysis (ICA; Beckmann and Smith, 2004)⍰, that also depend on an optimisation procedure. But to draw meaningful relations to behaviour, we need estimates that are robust and reproducible.

We introduce two approaches to enhance the stability in time-varying FC estimates. We focus on the HMM, but analogous solutions could be devised for other models. The first approach, termed best-ranked HMM (BR-HMM), consists of running the model several times and taking the run that best scores with respect to a given quantitative measure of model performance. In the case of the HMM, the natural choice is the free energy, which, based on Bayesian principles, represents a trade-off between fitness and model complexity. Previous studies have used the strategy of minimizing free energy to select the best model, typically considering only 10 to 20 runs (#refs). However, we demonstrate that, depending on the characteristics of the data, a larger number of runs might be needed to find a robust and reproducible model. The second approach, named hierarchical-clustered HMM (HC-HMM), involves using hierarchical clustering to group the state timeseries obtained from multiple HMM runs based on their similarities. The states within each cluster are then aggregated to create a more robust representation of time-varying FC. We show the effectiveness of these methods on two separate fMRI and magnetoencephalography (MEG) datasets. Our results demonstrate that HC-HMM is more computationally efficient than BR-HMM when the variability in the model inference is high. However, when the variability is low, BR-HMM outperforms HC-HMM.

## Materials and Methods

### Data description

#### Resting-state fMRI HCP data

We used publicly available fMRI data from 100 subjects from the HCP (Van Essen et al., 2013)□. For each participant, we considered data from 4 resting-state sessions of approximately 15 min each. Please refer to (Van Essen et al., 2012)□ for full details about the acquisition and preprocessing of the data. In brief, 3T whole-brain fMRI data were acquired with a spatial resolution of 2×2×2 mm and a temporal resolution of 0.72 s. All fMRI processing was performed using FSL (Jenkinson et al., 2012)□ including minimal high-pass temporal filtering (>2000s FWHM) to remove the linear trends of the data, and artefact removal using ICA+FIX (Griffanti et al., 2014)□. No low-pass temporal filtering or global signal regression was applied. Group spatial ICA was performed using MELODIC (Beckmann and Smith, 2004)□ to obtain a parcellation of 25 ICs. These timeseries were directly obtained from the HCP ‘PTN’ Parcellation+Timeseries+Netmats (specifically, the first 100 subjects from the ‘recon2’ version). Whereas the HCP provides a higher number of IC parcellations, the 25-ICA parcellation was sufficient to map the dynamics of FC as it covers the major functional networks, while keeping the computational cost low. The number of rows (= 1200 x 4 x 100) in this dataset matrix equals the number of time points per session and subject and the number of columns (= 25) corresponds to the number of IC components.

#### Resting-state MEG UK-MEG data

The MEG dataset consisted of 5-min resting-state recordings from 10 subjects obtained from the UK MEG Partnership (Hunt et al., 2016)□ which recruited 77 healthy participants at the University of Nottingham. Resting state MEG data were acquired using a 275-channel CTF MEG system (MISL, Coquitlam, Canada) operating in third-order synthetic gradiometry configuration, at a sampling frequency of 1200⍰ZHz. For MEG coregistration, MRI data collected with a Phillips Achieva 7T was employed. MEG data were downsampled to 250 Hz using an anti-aliasing filter, filtered out frequencies <1 Hz, and source-reconstructed to 42 dipoles (covering whole-brain cortical but not subcortical regions) using a linearly constrained minimum variance scalar beamformer. Out of these dipoles, 38 were created through ICA decomposition on resting-state fMRI data from the HCP, which was previously used to estimate large-scale static FC networks in MEG (Colclough et al., 2016). The remaining dipoles relate to the anterior and posterior precuneus. A weighted mask was employed to project the data to brain space, with each area having its highest value near the centre of gravity. Bad segments were manually eliminated, and spatial leakage was corrected using the technique described in Colclough et al. (2015). Twenty-two subjects were excluded due to excessive head motion or artefacts. From the remaining 55 participants (mean age 26.5 years, maximum age 48 years, minimum age 18 years, 35 men), the current study employed a selection of 10 subjects. This number was chosen to reduce the computational cost of our analysis. The number of rows (= 725006) in this dataset matrix corresponds to the number of time points across subjects’ sessions, and the number of columns (= 42) corresponds to the number of brain regions.

### Analysis pipeline

We applied the following methods on each dataset: (i) HMM, (ii) BR-HMM and (iii) HC-HMM. The results of the fMRI and the MEG datasets are presented in the same order.

### Hidden Markov Model (HMM)

Our analysis begins by estimating time-varying FC patterns using an HMM. The model assumes that timeseries can be described using a hidden sequence of a finite number of states. In a data-driven way, the HMM characterizes neural timeseries of concatenated data (**Figure 1a**) using a finite number of *K* (group-level) states that reoccur across time (**Figure 1b**). Each state timeseries represents the probability for that state to be active at each time point. Note that the spatial parameters of the HMM are at the group level, while the state timeseries are specific to each subject. The states are probability distributions within a certain family, and each of them is characterized by a certain set of parameters. To focus on FC, each state was here characterized by a Gaussian distribution with no mean parameter and a full covariance matrix representing pairwise covariation across regions (equivalent to using a Wishart distribution). For fMRI data, this was run on the BOLD signal, whereas for MEG this was run on the power (frequency filtered into the alpha-band; 8–12 Hz). Setting the mean vector of all HMM states to zero can be used to avoid confounding effects of mean activity level changes on FC patterns (Vidaurre et al., 2021)□□. Finally, a K-by-K transition probability matrix is estimated as part of the model, containing the probability of transitioning from one state to another or remaining in the current state. As is typical in HMM analysis, the transition probability matrix and state distribution parameters were estimated using the Baum-Welch algorithm, while the forward-backward algorithm was used to estimate the state timeseries.

**Figure 1.**
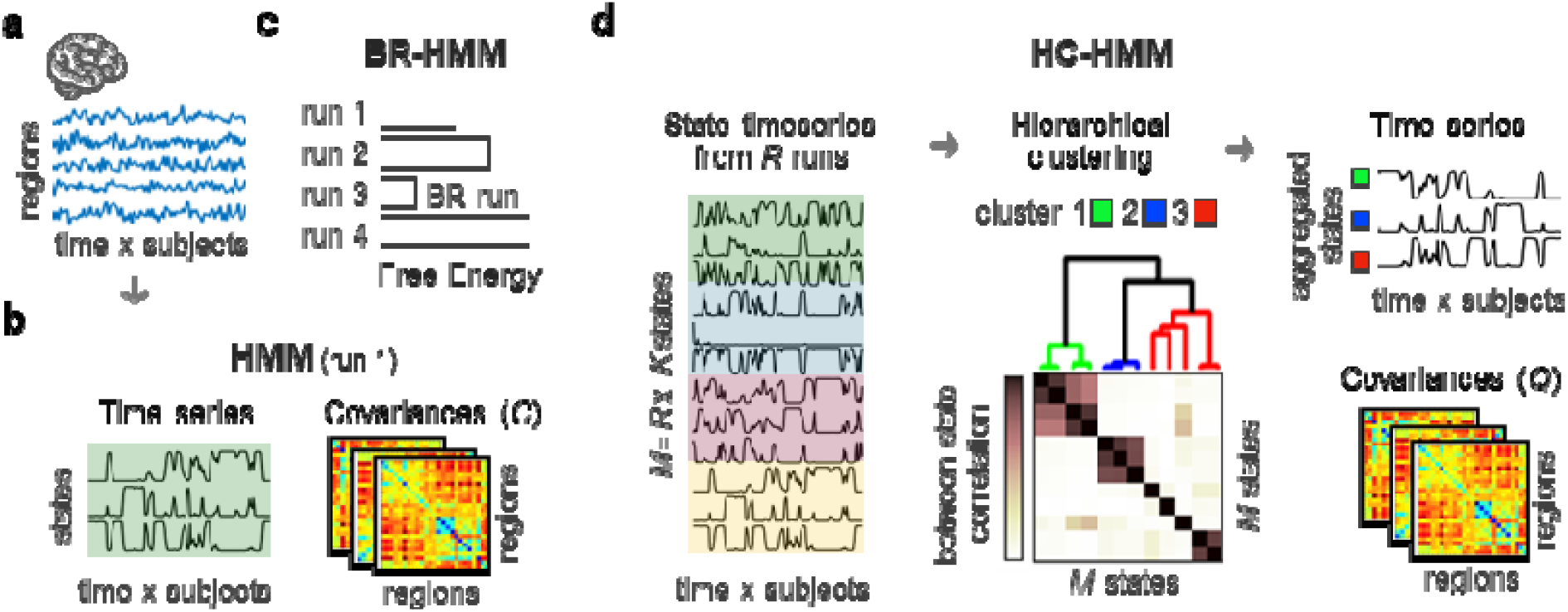
Graphical summary of the two approaches (BR-HMM and HC-HMM) to achieve reliable patterns of time-varying FC. **a)** fMRI or MEG timeseries, temporally concatenated across subjects. **b)** An HMM with K states was inferred from the concatenated brain timeseries to obtain patterns of time-varying FC (i.e., covariance matrices across regions). For the MEG dataset, the HMM was run on power timeseries. The state timeseries contain the probability of a given state to be active at each time point. **c)** BR-HMM approach consists of running the model inference multiple times, each starting from a different random initialization, and selecting the best-ranked HMM run according to the free energy, which is a quantitative measure that weighs the fitness of the data and simplicity of the model. **d**) HC-HMM approach involves running the HMM multiple times (*R*) and clustering the resulting state timeseries (*M=R×K*) according to their Pearson’s correlation to obtain more stable time-varying FC estimates. Ward’s hierarchical clustering algorithm is then applied to the *M×M* between-state correlation matrix to find similar state clusters. By averaging the original HMM state timeseries within each cluster, a set of aggregated state timeseries are produced. The covariances of each aggregated state are calculated by computing the weighted average of the covariance matrices of the original states within each cluster, where the weight assigned to each original state corresponds to its fractional occupancy, which represents the average probability of the state activation throughout the dataset. BR-HMM: Best-ranked HMM; HC-HMM: Hierarchical-clustered HMM; HMM: hidden Markov model(ling).

The HMM relies on an optimization procedure that starts from a random initialization point. This means that the final model parameters and results can be sensitive to the initial conditions and can vary from one run to another, even on the same dataset. This sensitivity to the initial conditions is a source of statistical noise in HMMs which can lead to less stable results. To evaluate how much the results across runs differed from each other, we run the model multiple times and compared the similarity of the estimates across runs. The *similarity* between two runs was measured as the statistical dependence between two sets of state timeseries. Because the ordering of the states within a run is arbitrary, the states of a run were first aligned to the states of the other run using the Hungarian algorithm (Munkres, 1957; Vidaurre, 2021)□: an optimization method that matches the states of two runs by minimising a cost matrix that reflects the dissimilarity between the aligned states.

### Best-ranked Hidden Markov Model (BR-HMM)

In the BR-HMM approach (**Figure 1c**), we run the model multiple times and choose the run with the lowest free energy. The free energy in the HMM inference is a Bayesian measure of a trade-off between model fitness to the data and complexity (given by how the posteriors differ from the priors). The mathematical formulas for calculating free energy in relation to HMM can be found in another source (Vidaurre et al., 2016)□.

To determine the minimum number of HMM runs required for the BR-HMM approach to produce stable results, we varied the number of HMM runs (*R*) from 5 to 500 and performed eight BR-HMM repetitions for each value of *R*. That is, in each repetition, the HMM was run *R* times, and the run with the lowest free energy was chosen as the BR-HMM run. We then assessed the correspondence across estimates by measuring the similarity between each pair of BR-HMM runs.

### Hierarchical-clustered Hidden Markov Model (HC-HMM)

The HC-HMM approach (**Figure 1d**) uses a hierarchical clustering to group the *K* state timeseries obtained from multiple runs of an HMM. The generated clusters are used to produce more stable state timeseries by aggregating the original HMM state timeseries within each cluster. This aggregation process integrates the multiple solutions obtained across HMM runs, resulting in a more reliable representation of the underlying patterns of time-varying FC.

Specifically, a hierarchical clustering algorithm was fed the between-state correlation matrix, *P*. This matrix was created by calculating the Pearson correlation between every pair of state timeseries. The total number of states is *M=K×R*. As a distance measure, we used *D*_ij_=1−*P*_*ij*_, where *P*_*ij*_ represents the correlation coefficient between the i^th^ and j^th^ state. The clustering algorithm starts by regarding each element as a separate cluster; then the closest pair of clusters are iteratively combined into a larger cluster. To measure the distance between two clusters, we used Ward’s linkage, which minimizes the variance of the clusters being merged. As a result, highly correlated states will be clustered together. A dendrogram was obtained from this procedure; and each of the clusters can be seen as a group of states with the same underlying FC pattern. We set the number of clusters to match the number of states (*K*) in the HMM. The resulting clusters were then used to compute aggregated state timeseries by averaging the timeseries within each cluster. To estimate the spatial patterns of time-varying FC associated with each aggregated state timeseries, we computed one covariance matrix for each aggregated state (denoted as *Q*_*i*_) by taking the weighted average of the covariance matrices of the original states (*C*_*m*_) within that cluster, where the weights correspond to the fractional occupancy of each original state, i.e., the average probability of a state activation throughout the dataset.

Again here, to evaluate the stability of the results, HC-HMM was repeated eight times and the similarity of the results across repetitions were compared. We also quantified the relationship between HC-HMM stability and the number of HMM runs, for various choices of *R* (from 5 to 300).

### Computational resources and environment

The computations were performed using MATLAB version 9.10 (R2021a) on an AMD Ryzen 9 3900X 12-Core CPU running the Linux 5.15.0-52-generic operating system. The system had 52 GB of RAM available. A single run of the HMM on the HCP dataset took a total of 477.44 seconds (∼7.96 minutes) to complete. For the MEG dataset one run took 118.60 seconds (∼1.98 minutes) to complete.

## Results

The following results, for both fMRI and MEG, follow the analysis pipelines detailed in the *Materials and Methods* section. All models were run at the group level, and all results are group-level results.

### FMRI data

The fMRI dataset considered here consisted of four sessions of resting-state fMRI data from 100 subjects obtained from the HCP (Van Essen et al., 2013)□. The data were projected onto 25 independent components (ICs), which we used as input timeseries. Whole-brain patterns of time-varying FC were obtained by applying an HMM to the concatenated (standardized) timeseries for all subjects (25 ICs by [time × sessions × subjects]). Here the HMM was modelled with *K*=12 states, where each state represents a reoccurring pattern of FC.

### HMM stability

The stability of the HMM inference was evaluated using the between-run similarities across 1000 runs. The similarity score has a value between 0 and 1, which is proportional to how close the set of states of a run is to the set of states of a different run. A histogram of the similarity scores for each pair of runs revealed a minimum similarity score of 0.36, a maximum of 0.99, and an average score of 0.59 (**Figure 2a)**. This indicates that the states inferred by the HMM are to some extent different across runs of the inference, as shown previously in Vidaurre et al. (2019).

**Figure 2.**
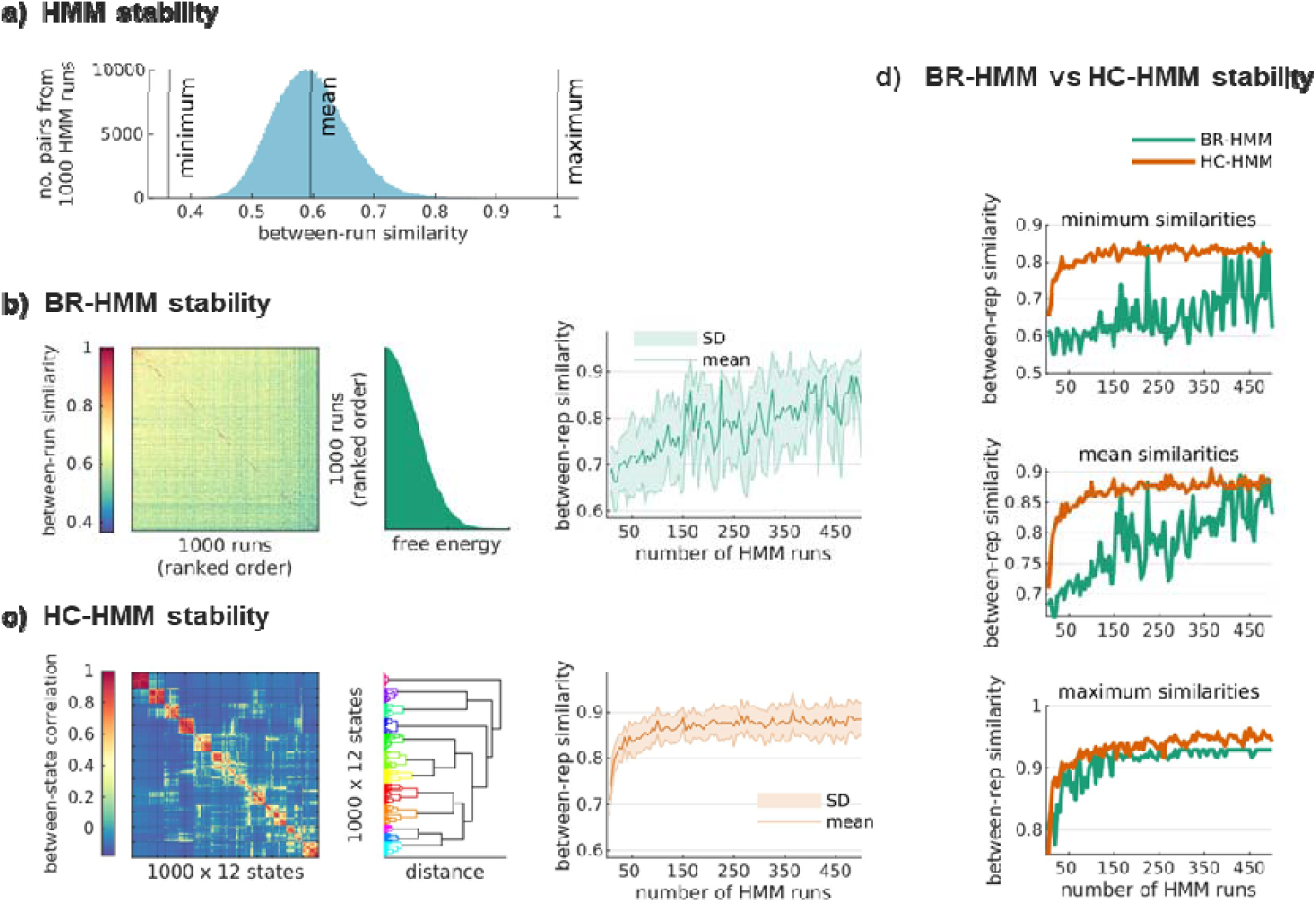
Capturing more stable time-varying FC estimations from resting-state fMRI data. **a**) Histogram of between-run similarities computed from 1000 HMM runs (N=499500 pairs of runs). The similarity of each pair of HMM runs was computed as the sum of the joint probabilities across two sets of *K*=12 state timeseries (*see Materials and Methods*). **b)** (*left* to *right*) Run-by-run matrix of the between-run similarities, with the runs sorted in ascending order based on their free energy; free energy levels for each HMM run sorted in ascending order; between-repetition similarities of the BR-HMM approach as a function of the number of HMM runs *R*, from 5 to 500, in steps of 5. **c)** (*left* to *right*) Matrix of Pearson correlation coefficients between pairs of state timeseries with states ordered according to hierarchical clustering (*K* clusters, the total number of states is *R×K* =12000); dendrogram showing the states within each cluster; between-repetition similarities of the HC-HMM approach as a function of *R* (from 5 to 500, in steps of 5). **d)** (*left* to *right*) Overlay plots of the minimum, mean and maximum similarity scores across repetitions as a function of *R* (BR-HMMs in green, and HC-HMMs in orange).

### BR-HMM stability

In the BR-HMM approach, we chose the run that best-ranked according to the free energy. How well this approach will work depends on whether the estimates generated from BR-HMM runs (i.e., the runs with the lowest free energy) all represent the same underlying HMM solution. In **Figure 2b**-*left*, a matrix of similarities is presented across the estimates obtained from 1000 HMM runs, sorted in ascending order based on their free energy values. The between-run similarity matrix reveals that the HMM runs with the lowest free energy values are very similar to each other.

To better evaluate the stability of BR-HMM runs, which are defined as the HMM runs with the lowest free energy among *R* runs, we conducted eight repetitions of the BR-HMM approach. We then assessed the similarity between the results across repetitions for different numbers of HMM runs, ranging from *R*=5 to *R*=500. The stability of the BR-HMM estimates gradually improves as the number of runs increases, as depicted in **Figure 2b**-*right*. However, the standard deviation of approximately 0.08 indicates that, still, there is some variability across BR-HMM runs. While some pairs of BR-HMM runs have a similarity of 0.93, others cannot reach 0.7, suggesting that not all BR-runs converge to the same underlying HMM solution. Therefore, many HMM runs (here, 500) might be needed for reaching a robust solution.

Overall, our findings suggest that the stability of BR-HMM estimates is dependent on having enough runs of the HMM to adequately describe the landscape of possible solutions. As the complexity of the model increases, more runs are needed, which can be computationally intensive.

### HC-HMM stability

The HC-HMM approach utilizes hierarchical clustering to group state timeseries obtained from multiple HMM runs based on their similarity. The clustering is based on a between-state similarity matrix, given by correlation between the state time series. The total number of states was *R×K*=1000×12; the number of clusters was set to 12 to match *K*. In **Figure 2c**-*left*, a matrix of the Pearson correlation coefficients between each pair of state timeseries is presented, with states ordered based on the hierarchical clustering to visualize clusters of similar states. Clusters of similar states tend to concentrate around the main diagonal. It is important to note that lack of correlation between clusters is expected as the states derived from the HMM are mutually exclusive (i.e., the sum of the probabilities of activation across states at a given time point for a given HMM is 1). The resulting clusters are depicted in the dendrogram of **Figure 2c**-*middle* and were used to generate aggregated timeseries by averaging the timeseries within each cluster. This method provides a robust and stable estimation of state timeseries by integrating information from multiple runs of the HMM.

To determine the minimum number of runs required for the HC-HMM approach to return stable results, we assessed its performance across various numbers of runs, ranging from 5 to 500. To do so, we repeated the HC-HMM procedure eight times for each value of R and compared the similarities across the aggregated state timeseries (**Figure 2c**-*right*). Our results showed that the HC-HMM approach achieved prominent levels of stability with just 50 runs, with similarities across HC-HMM estimates exceeding 0.84. In general, adding more runs did not significantly improve the stability of HC-HMM beyond 0.89 (SD = 0.03).

### BR-HMM versus HC-HMM stability

The between-repetition similarities of both approaches are also represented in **Figure 2d** in an overlay plot to facilitate comparisons of the two methods. The minimum, mean, and maximum similarity scores across repetitions of HC-HMM were consistently higher than the BR-HMM. The HC-HMM approach achieved stability above 0.84 with just 50 runs, while more than 500 runs were needed for BR-HMM to reach the same levels of stability. Further details about the underlying FC patterns can be found in **Supplementary Figure 1**.

Taken together, despite the stochastic nature of the HMM inference, momentary changes in FC can be reliably captured in fMRI. However, while the BR-HMM approach required hundreds of inference realizations to describe the landscape of possible HMM solutions, HC-HMM achieved high stability with just a fraction of the runs. This highlights the advantages of the HC-HMM approach, particularly when inference is variable enough that the number of runs needed to obtain minimal values of free energy is high.

### MEG data

The MEG dataset considered here consists of 5-minute resting-state recordings from 10 subjects. The data were mapped to a 42-region parcellation. HMMs with K=6 states were inferred from the temporally concatenated data across subjects to obtain whole-brain patterns of time-varying FC. For easier comparison with the fMRI results, the HMM was applied on the power (band-limited to 8–12 Hz) instead of the raw timeseries; *see Materials and Methods* for further details.

### HMM stability

To evaluate the stability of the HMM inference for a model with *K*=6 states, the model was run 1000 times and the similarity across runs was computed. **Figure 3a** displays a histogram of the similarities across runs, which shows a minimum similarity score of 0.37, a maximum of 0.97, and an average score of 0.65 (SD=0.09). These results indicate a slightly lower inference variability compared to the fMRI dataset.

**Figure 3.**
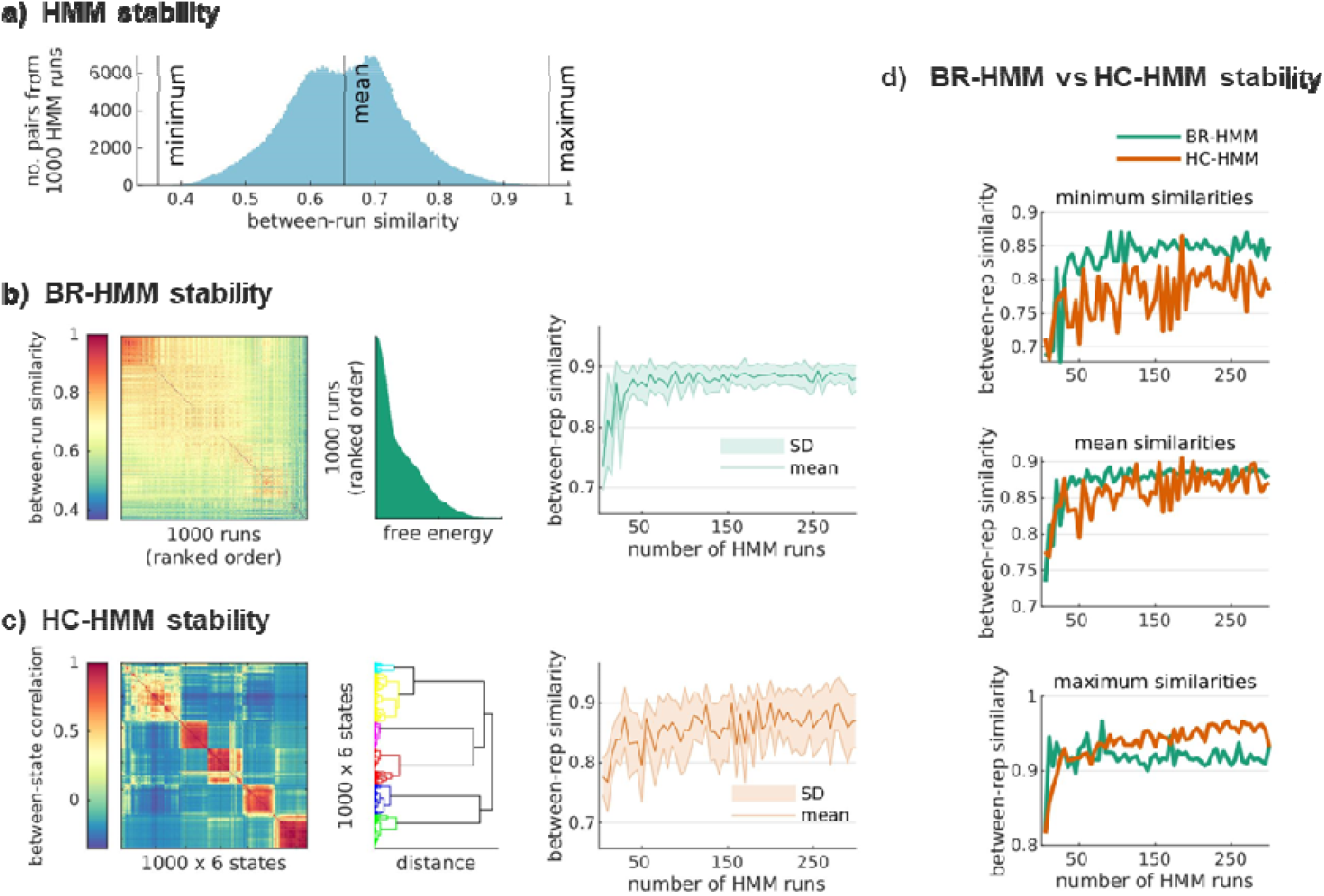
Capturing more stable time-varying FC estimations from resting-state (alpha-band) MEG data. **a)** Histogram of between-run similarities computed from 1000 HMM runs (N=499500 pairs of runs). The similarity of each pair of HMM runs was computed as the sum of the joint probabilities across two sets of *K*=6 state timeseries (*see Materials and Methods*). **b)** (*left* to *right*) Run-by-run matrix of the between-run similarities, with the runs sorted in ascending order based on their free energy; free energy levels for each HMM run sorted in ascending order; between-repetition similarities of the BR-HMM approach as a function of the number of HMM runs *R*, from 5 to 500, in steps of 5. **c)** (*left to right*) Matrix of Pearson correlation coefficients between pairs of state timeseries with states ordered according to hierarchical clustering (*K* clusters, the total number of states is *R×K* =6000); dendrogram showing the states within each cluster; between-repetition similarities of the HC-HMM approach as a function of *R* (from 5 to 500, in steps of 5). **d**) (*left* to *right*) Overlay plots of the minimum, mean and maximum similarity scores across repetitions as a function of R (BR-HMMs in green, and HC-HMMs in orange).

### BR-HMM stability

Like in the fMRI dataset, the BR-HMM approach was used to select the best run among multiple HMM runs based on the lowest free energy value. **Figure 3b**-*left* shows the free energy values for the 1000 HMM runs sorted in ascending order, with highly stable runs having the lowest free energy values, as evidenced by the high similarity scores among these runs. The relationship between free energy and stability in this dataset was clearer than in the fMRI dataset.

To determine the number of HMM runs needed to reach a stable minimal value of free energy, the stability of estimates obtained from BR-HMM runs was evaluated through eight repetitions of the BR-HMM method. The similarity of results across repetitions was compared for different numbers of HMM runs, ranging from *R*=5 to *R*=500. As the number of HMM runs increased, the stability of estimates obtained from BR-HMM runs improved considerably, with a stable mean similarity of 0.87 and a standard deviation of 0.06 reached at around 50 runs, as shown in **Figure 3b**-*right*. Adding more runs did not lead to significant improvements in stability beyond this point.

### HC-HMM stability

After running multiple HMM runs, we used the HC-HMM approach to group state timeseries based on their similarity. As before, the clustering was performed on a between-state correlation matrix that captured the correlation between each pair of state timeseries. Pearson correlation coefficients between each pair of state timeseries are shown in **Figure 3c**-*left*, with the states ordered based on the hierarchical clustering. The resulting clusters were depicted in a dendrogram; aggregated timeseries were generated by averaging the timeseries within each cluster.

To assess the stability of the HC-HMM approach, we repeated the procedure eight times for each value of *R* and compared the similarities across the aggregated state timeseries for various numbers of runs (5 to 500). The results showed that the HC-HMM approach consistently yielded higher similarities between aggregated states obtained from the original HMM states over as few as 20 runs (**Figure 3c-***right*).

### BR-HMM versus HC-HMM stability

The stability trends obtained using the BR-HMM and HC-HMM were similar. While the mean stability of BR-HMM is only slightly better than HC-HMM, the latter shows a wider standard deviation. Therefore, although the estimations across repetitions of the HC-HMM approach may reach values close to one, or even higher than those of the BR-HMM approach, this is not always the case.

Taken together, our results demonstrate that both approaches can improve the stability of time-varying FC patterns in MEG data. However, when the lowest free-energy runs represent varying forms of the same HMM decomposition, then, the simpler BR-HMM approach may be sufficient to capture the underlying dynamics, and the additional complexity introduced by HC-HMM may hinder its performance. Further details about the underlying FC patterns can be found in **Supplementary Figure 2**.

## Discussion

Inference variability from complex models like the HMM means that different runs of the inference algorithm can produce different results. This can lead to decreased reproducibility and interpretability of results, which in turn can impact our ability to make accurate predictions or identify meaningful relationships with behaviour. Additionally, time-varying FC is a fundamental component of many dynamic metrics used in network neuroscience, such as graph measures and network properties, which quantify the topology and organization of brain networks and their changes over time. Variability in the estimation of time-varying FC can lead to unstable and unreliable dynamic metrics, affecting the interpretation and validity of the results.

Many studies in the field of network neuroscience use sliding windows to estimate time-varying FC. The sliding window approach involves dividing the timeseries data into smaller time windows and estimating the FC for each window. By sliding the window across the timeseries, the FC can be estimated as a function of time. However, this approach has limitations because the estimation of FC is inherently noisy and can be affected by variations in the length and placement of the windows, resulting in a high degree of statistical variability. The hidden Markov model (HMM) is an alternative that, by using a finite number of states to characterize time-varying FC (Vidaurre et al., 2017)□□, does not suffer from such limitations and allows modelling time point by time point variability of FC. The inference of model parameters from the data in the HMM (as in many other models) relies on an optimisation procedure. Due to the stochastic nature of the inference together with the inherent noise in the data, different runs of the inference algorithm can produce different estimates of the model parameters, i.e., different estimates of time-varying FC. In this study, we proposed two approaches, BR-HMM and HC-HMM, to improve the stability of HMM estimates. Our results demonstrated that both methods can greatly improve the stability of the estimates.

The BR-HMM approach selects the best estimate of the time-varying FC by minimizing the free energy. While running the HMM multiple times can improve performance and reduce variability, the computational cost of the BR-HMM approach can be a limiting factor, particularly when the number of runs needed to obtain minimal values of free energy is high. This is especially true for high-complexity models, which require more runs to explore the space of possible solutions. In such cases, HC-HMM is recommended, as it is better suited to handle greater complexity of the data. Alternatives to hierarchical clustering would be the use of principal component analysis or non-negative matrix factorization. However, unlike the HC-HMM approach, these would produce components that do not follow the assumptions of the HMM and are therefore harder to interpret. Having an HMM-compatible model structure is particularly useful for characterising individual FC profiles, including their activation probability and duration, as well as identifying state transitions. A potential avenue for further improvement in the HC-HMM method is related to the selection of the number of clusters. In this study, we chose to match the number of clusters to the number of states in each individual HMM. However, this may not always be the most appropriate choice. A data-driven approach for determining the optimal number of clusters could potentially yield more stable results. One method would be the elbow criterion, which identifies the point where the rate of change in the sum of squares levels off. Another option would be to visually inspect the cluster assignments and exclude any states that do not clearly belong to any of the clusters, which could improve the interpretability and robustness of the results obtained from the HC-HMM.

Finally, it is important to consider that the proposed methods for reducing the inference variability of the HMM address the instability that arises from the randomness of the optimization procedure used to estimate the model parameters. However, there may be other sources of instability in the estimation process that are not addressed by these methods. For example, the complexity of the landscape of FC dynamics may make it difficult to accurately describe the data with the parameters of the model and this could contribute to variability in the estimation results. Alternatively, estimation noise may arise due to limitations in the amount or quality of data available or high signal-to-noise ratio probabilities (Lee et al., 2010)L. To further improve the accuracy and stability of the model, future work may need to focus on reducing the variability due to these other factors. One approach could involve varying the model parameters in combination with the proposed methods for reducing variability due to optimization randomness. By fine-tuning the model parameters to better capture the complexity of the FC dynamics or accounting for estimation noise, it may be possible to further reduce the variability in the estimation results and obtain more reliable and accurate estimates of time-varying FC.

## Conclusion

Reproducibility is a growing concern in neuroimaging, and this issue can be further complicated by the stochastic nature of the different methods’ inference. In this study, we have focused on the hidden Markov model to characterising time-varying functional connectivity and have proposed two methods to achieve stable estimations: BR-HMM and HC-HMM. These methods significantly improved the stability of the estimations up to almost 0.9. While BR-HMM, can produce higher stability scores, its computational cost may be a limiting factor in some cases. On the other hand, HC-HMM offers a computationally affordable solution. By reducing the inference variability that originates from the randomness of the optimization procedure, the proposed methods can yield more stable and reliable estimates of time-varying FC, which can aid in our understanding of neural processes and cognitive functions.

## Data Availability

The fMRI data were provided by the Human Connectome Project, WU-Minn Consortium (Principal Investigators: David Van Essen and Kamil Ugurbil; 1U54MH091657) funded by the 16 NIH Institutes and Centers that support the NIH Blueprint for Neuroscience Research; and by the McDonnell Center for Systems Neuroscience at Washington University. The MEG dataset is part of a larger dataset acquired in Nottingham in the context of the MEG UK Partnership and is not currently available as it contains data from human participants including structural scans. The data are held by the MEG UK Partnership, and access to the MEG UK Database can be requested at http://meguk.ac.uk/contact. Preprocessed (and parcellated) data containing the time series as they were fed to the HMM can be accessed at https://github.com/sonsolesalonsomartinez/reproducibleHMM.

## Code Availability

The software comprises the open-access code repositories https://github.com/OHBA-analysis/HMM-MAR and custom Matlab scripts https://github.com/sonsolesalonsomartinez/reproducibleHMM.

## Acknowledgments

DV is supported by a Novo Nordisk Foundation Emerging Investigator Fellowship (NNF19OC-0054895) and an ERC Starting Grant (ERC-StG-2019–850404). This research was funded in part by the Wellcome Trust (215573/Z/19/Z). For the purpose of Open Access, the authors have applied a CC BY NC ND public copyright licence to any Author Accepted Manuscript version arising from this submission.

## References

Baker AP, Brookes MJ, Rezek IA, Smith SM, Behrens T, Probert Smith PJ, Woolrich M. 2014. Fast transient networks in spontaneous human brain activity. Elife 3: e01867. doi: 10.7554/eLife.01867.

Beckmann CF, Smith SM. 2004. Probabilistic Independent Component Analysis for Functional Magnetic Resonance Imaging. IEEE Trans Med Imaging 23:137–152. doi:10.1109/TMI.2003.822821

Bellaïche A. 1996. The tangent space in sub-Riemannian geometry. Sub-Riemannian Geom 1–78. doi:10.1007/978-3-0348-9210-0_1

Colclough GL, Brookes MJ, Smith SM, Woolrich MW. A symmetric multivariate leakage correction for MEG connectomes. Neuroimage 117, 439–448 (2015).

Colclough GL, Woolrich MW, Tewarie PK, Brookes MK, Quinn AJ, Smith SM. 2016. How reliable are MEG resting-state connectivity metrics? Neuroimage 138, 284–293.

Congedo M, Barachant A, Bhatia R. 2017. Riemannian geometry for EEG-based brain-computer interfaces; a primer and a review. http://dx.doi.org/101080/2326263X20171297192 4:155–174. doi:10.1080/2326263X.2017.1297192

Dafflon J, F. Da Costa P, Váša F, Monti RP, Bzdok D, Hellyer PJ, Turkheimer F, Smallwood J, Jones E, Leech R. 2022. A guided multiverse study of neuroimaging analyses. Nat Commun 2022 131 13:1–13. doi:10.1038/s41467-022-31347-8

de Pasquale F, Della Penna S, Snyder AZ, Lewis C, Mantini D, Marzetti L, Belardinelli P, Ciancetta L, Pizzella V, Romani GL, Corbetta M. 2010. Temporal dynamics of spontaneous MEG activity in brain networks. Proc Natl Acad Sci U S A. 107:6040–6045.

de Pasquale F, Della Penna S, Snyder AZ, Marzetti L, Pizzella V, Romani GL, Corbetta M. 2012. A cortical core for dynamic integration of functional networks in the resting human brain. Neuron 74(4):753–64. doi: 10.1016/j.neuron.2012.03.031.

Fornito A, Bullmore ET. 2010. What can spontaneous fluctuations of the blood oxygenation-level-dependent signal tell us about psychiatric disorders? Curr Opin Psychiatry 23:239–49. doi:10.1097/YCO.0b013e328337d78d

Friston KJ. 1994. Functional and effective connectivity in neuroimaging: A synthesis. Hum Brain Mapp 2:56–78. doi:10.1002/hbm.460020107

Griffanti L, Salimi-Khorshidi G, Beckmann CF, Auerbach EJ, Douaud G, Sexton CE, Zsoldos E, Ebmeier KP, Filippini N, Mackay CE, Moeller S, Xu J, Yacoub E, Baselli G, Ugurbil K, Miller KL, Smith SM. 2014. ICA-based artefact removal and accelerated fMRI acquisition for improved resting state network imaging. Neuroimage 95:232–247. doi: 10.1016/j.neuroimage.2014.03.034

Harrison SJ, Woolrich MW, Robinson EC, Glasser MF, Beckmann CF, Jenkinson M, Smith SM. 2015. Large-scale probabilistic functional modes from resting state fMRI. Neuroimage 109:217–231. doi: 10.1016/j.neuroimage.2015.01.013

Hindriks R, Adhikari MHH, Murayama Y, Ganzetti M, Mantini D, Logothetis NKK, Deco G. 2016. Can sliding-window correlations reveal dynamic functional connectivity in resting-state fMRI? Neuroimage 127:242–256. doi: 10.1016/j.neuroimage.2015.11.055

Hunt BAE, Tewarie PK, Mougin OE, Geades N, Jones DK, Singh KD, Morris PG, Gowland PA, Brookes MJ. 2016. Relationships between cortical myeloarchitecture and electrophysiological networks 113:13510–13515.

Jenkinson M, Beckmann CF, Behrens TEJ, Woolrich MW, Smith SM. 2012. FSL. Neuroimage 62:782–790. doi: 10.1016/j.neuroimage.2011.09.015

Karapanagiotidis T, Vidaurre D, Quinn AJ, Vatansever D, Poerio GL, Turnbull A, Ho NSP, Leech R, Bernhardt BC, Jefferies E, Margulies DS, Nichols TE, Woolrich MW, Smallwood J. 2020. The psychological correlates of distinct neural states occurring during wakeful rest. Sci Rep 10:21121. doi:10.1038/s41598-020-77336-z

Laughlin SB, Sejnowski TJ. 2003. Communication in Neuronal Networks. Science (80-) 301:1870–1874. doi:10.1126/science.1089662

Lee S, Huang JZ, Hu J. 2010. Sparse logistic principal components analysis for binary data. Ann Appl Stat 4:1579–1601. doi:10.1214/10-AOAS327

Liégeois R, Li J, Kong R, Orban C, Van De Ville D, Ge T, Sabuncu MR, Yeo BTT. 2019. Resting brain dynamics at different timescales capture distinct aspects of human behavior. Nat Commun 10:2317. doi:10.1038/s41467-019-10317-7

Lurie DJ, Kessler D, Bassett DS, Betzel RF, Breakspear M, Keilholz S, Kucyi A, Liégeois R, Lindquist MA, McIntosh AR, Poldrack RA, Shine JM, Thompson WH, Bielczyk NZ, Douw L, Kraft D, Miller RL, Muthuraman M, Pasquini L, Razi A, Vidaurre D, Xie H, Calhoun VD. 2020. Questions and controversies in the study of time-varying functional connectivity in resting fMRI. Netw Neurosci 4:30–69. doi:10.1162/NETN_A_00116

Munkres J. 1957. Algorithms for the Assignment and Transportation Problems. J Soc Ind Appl Math 5:32–38. doi:10.1137/0105003

Van Essen DC, Smith SM, Barch DM, Behrens TEJ, Yacoub E, Ugurbil K. 2013. The WU-Minn Human Connectome Project: An overview. Neuroimage 80:62–79. doi: 10.1016/j.neuroimage.2013.05.041

Van Essen DC, Ugurbil K, Auerbach E, Barch D, Behrens TEJ, Bucholz R, Chang A, Chen L, Corbetta M, Curtiss SW, Della Penna S, Feinberg D, Glasser MF, Harel N, Heath AC, Larson-Prior L, Marcus D, Michalareas G, Moeller S, Oostenveld R, Petersen SE, Prior F, Schlaggar BL, Smith SM, Snyder AZ, Xu J, Yacoub E. 2012. The Human Connectome Project: A data acquisition perspective. Neuroimage 62:2222–2231. doi: 10.1016/j.neuroimage.2012.02.018

Vidaurre D. 2021. A new model for simultaneous dimensionality reduction and time-varying functional connectivity estimation. PLOS Comput Biol 17(4): e1008580

Vidaurre D, Hunt LT, Quinn AJ, Hunt BAEE, Brookes MJ, Nobre AC, Woolrich MW. 2018. Spontaneous cortical activity transiently organises into frequency specific phase-coupling networks. Nat Commun 9:2987. doi:10.1038/s41467-018-05316-z

Vidaurre D, Llera A, Smith SMM, Woolrich MWW. 2021. Behavioural relevance of spontaneous, transient brain network interactions in fMRI. Neuroimage 229:117713. doi: 10.1016/j.neuroimage.2020.117713

Vidaurre D, Quinn AJ, Baker AP, Dupret D, Tejero-Cantero A, Woolrich MW. 2016. Spectrally resolved fast transient brain states in electrophysiological data. Neuroimage 126:81–95. doi: 10.1016/j.neuroimage.2015.11.047

Vidaurre D, Smith SM, Woolrich MW. 2017. Brain network dynamics are hierarchically organized in time. Proc Natl Acad Sci 114:12827–12832. doi:10.1073/pnas.1705120114

Vidaurre D, Woolrich MW, Winkler AM, Karapanagiotidis T, Smallwood J, Nichols TE. 2019. Stable between-subject statistical inference from unstable within-subject functional connectivity estimates. Hum Brain Mapp 40:1234–1243. doi:10.1002/hbm.24442

Xie H, Gonzalez-Castillo J, Handwerker DA, Bandettini PA, Calhoun VD, Chen G, Damaraju E, Liu X, Mitra S. 2018. Time-varying whole-brain functional network connectivity coupled to task engagement. Netw Neurosci 3:49–66. doi:10.1162/netn_a_00051

