## Supplementary Figures for "Towards stability of dynamic FC estimates in neuroimaging and electrophysiology: solutions and limits"

### corresponding to

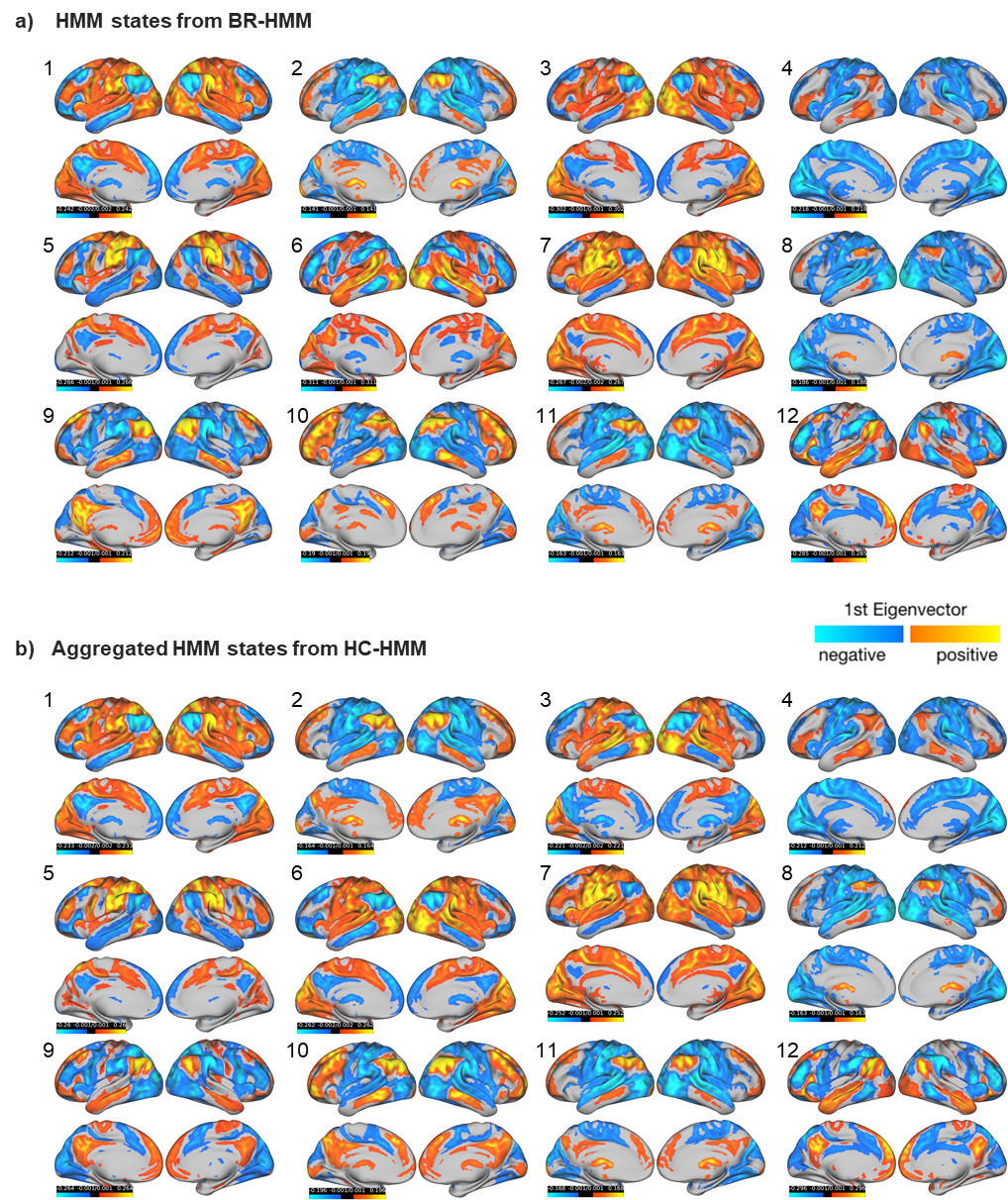

**Supplementary Figure 1. Spatial representation of fMRI FC states** derived from **a)** a BR-HMM and **b)** an HC-HMM. Brain connectivity maps were created using the 1st Eigenvector of each states’ covariance matrices.

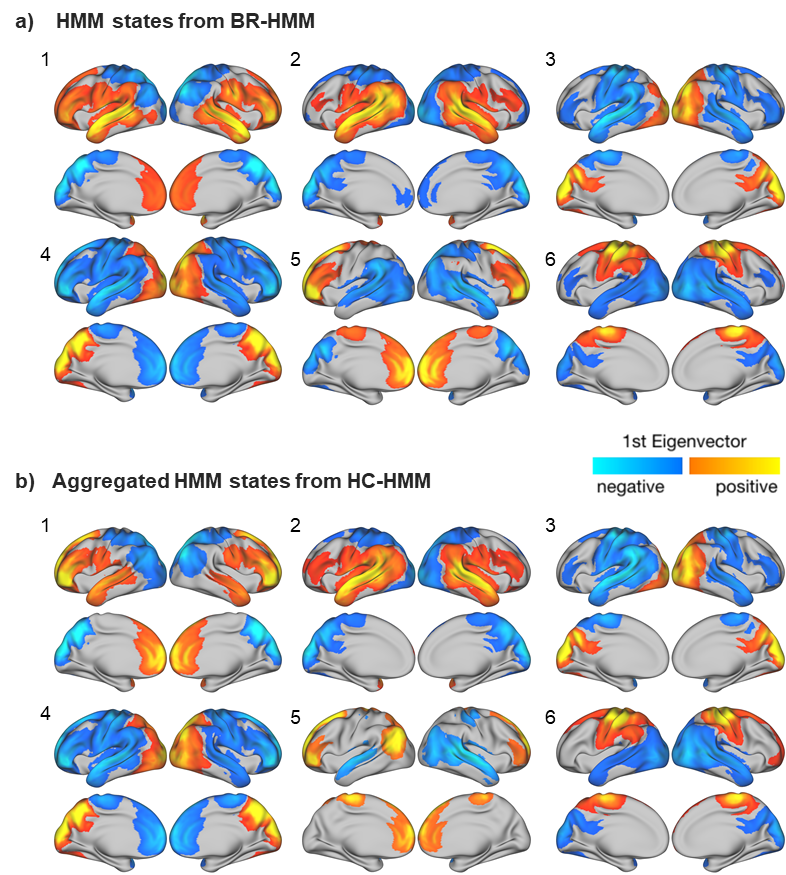

**Supplementary Figure 2. Spatial representation of MEG FC states** derived from **a)** a BR-HMM and **b)** an HC-HMM. Brain connectivity maps were created using the 1st Eigenvector of each state’s covariance matrices.
